# Avian diet and foraging ecology constrain foreign egg recognition and rejection

**DOI:** 10.1101/402941

**Authors:** Alec B. Luro, Mark E. Hauber

## Abstract

Egg rejection is the most common defense against avian brood parasitism in which the host either removes the parasitic egg or deserts the parasitized clutch. The ability to recognize and reject a parasitic egg depends on bill morphology, sensory systems, and cognition, all of which are also shaped by other selective processes, such as foraging. This begs the question whether specific phenotypes associated with different foraging strategies and diets may constrain or facilitate egg recognition and rejection. Here, we propose a novel hypothesis that host species phenotypes related to foraging ecology and diet impose morphological and sensory constraints on the evolution of egg rejection. We conducted a comparative analysis of the adult diets and egg rejection rates of 165 current host and non-host species and found species that consume an animal and fruit dominated diet rather than seeds and grains, forage arboreally rather than aerially, and possess relatively larger body sizes have significantly higher egg rejection rates. As predicted, these results suggest that phenotypes related to specific diets and foraging strategies may differentially constrain or facilitate evolution of host egg rejection defenses against avian brood parasitism.

## I. INTRODUCTION

Interspecific obligate avian brood parasitism is a breeding strategy wherein a parasitic species lays its eggs in the nests of other species that provide parental care for the unrelated offspring (Davies 2000). Egg rejection is the most common defense in which a host either removes the parasitic egg from the nest (Rothstein 1975) or deserts the parasitized nest (Hosoi & Rothstein 2000). The likelihood of hosts evolving egg rejection defense is predicted to be greater when the cost of raising brood parasite nestlings is higher (Medina & Langmore 2015). Yet, many host species do not reject foreign eggs, despite not only facing high costs of parasitism (Medina & Langmore 2015), but also having own eggs predicted to be readily perceived as visually different from brood parasite eggs (Stoddard & Stevens 2011), and demonstrating physical capability of removing objects from the nest as large and heavy as foreign eggs (Rothstein 1975).

There are several adaptive hypotheses for why some host species have not evolved egg rejection, including the evolutionary equilibrium hypothesis that suggests that the cost of breaking or mistakenly rejecting own eggs counterweigh the benefits of rejecting brood parasite eggs (Lotem et al. 1995). Alternatively, the nonadaptive evolutionary lag hypothesis proposes that not enough time has passed since the onset of a coevolutionary host-brood parasite arms race for the host to evolve egg rejection as a defense (Rothstein 1990).

A novel, non-exclusive hypothesis in concordance with evolutionary lag for the absence of host egg rejection is that different host species’ phenotypes may differentially constrain (Schwenk 1995) or facilitate the evolution of egg rejection over a period of time due to perceptual and physical limitations, regardless of the cost of raising unrelated offspring. Specifically, foreign egg recognition is only possible if the host is capable of perceiving brood parasite eggs as different from its own eggs, typically using visual cues (Stoddard & Hauber 2017). Furthermore, small host bill size or lower body mass may also limit the ability to remove a brood parasite egg from the nest (Rohwer & Spaw 1988).

How might species’ phenotypes have been shaped in a manner that either constrains or facilitates the evolution of egg rejection? Foraging ecology and diet play predominant roles in the evolution of avian body size, bill morphology, visual perception, sensory systems, and cognition (Mendelson et al. 2016; Pigot et al. 2016; Martin 2017), all of which may contribute to the ability to recognize and remove a foreign egg from the nest (Rohwer & Spaw 1988; Stoddard & Hauber 2017). For example, mostly granivorous, seed eating species generally have bills with wide depths and short lengths that may be unsuitable for piercing or grasping and removing foreign eggs (Rohwer & Spaw 1988), and they can have visual acuities that may be considered myopic in comparison to longer-billed visually guided insectivores which feed on highly mobile and/or often cryptic prey (Dolan & Fernández-Juricic 2010; Moore et al. 2015). Furthermore, mostly frugivorous birds may be more sensitive to color differences than granivorous birds, because chromatic differences are reliable long-distance cues for identifying ripe fruits (Cazetta et al. 2009), whereas achromatic features, including shape and pattern contrast, are likely more reliable cues in identifying cryptic seeds on the ground (Porter 2013). Therefore, we predict that mostly insectivorous and frugivorous birds may possess visual sensory and perceptual traits better suited than those of mostly granivorous birds for visually discriminating foreign eggs in their nests; potentially by discriminating egg maculation pattern differences, which may be constrained visual spatial resolution (i.e., acuity), or perceiving and attending to eggshell background color differences, which may be constrained by photoreceptor sensitivities and densities (Hart 2001; Cassey et al. 2008; Spottiswoode & Stevens 2010; Stoddard et al. 2014). Here, we test how foraging ecology and diet-related phenotypes may influence evolutionary trajectories of host egg rejection defenses using phylogenetic comparative methods.

## II. METHODS

### Data

We collected data from published avian egg rejection studies for as many species as we could find (N=174), using “egg rejection” as key words, then searched for published diet data for the respective species and generated a complete dataset of both egg rejection and categorized diet data, matched with a set of 100 phylogenetic hypotheses from birdtree.org (Jetz et al. 2012) for N=165 species (supplementary data). For each species, our complete dataset included: weighted average egg rejection rate (weighted by study experiment sample sizes), average body mass (g), categorical diet and forage zone (from del Hoyo et al. 2017), the associated obligate avian brood parasite taxon (*Cuculidae* and *Molothrus*) and whether the species is a known current host of its associated brood parasite, i.e., “host status”. (from Johnsgard 1997 and Ortega 1998). Diet categories included: “frugivore” (N=3), “granivore” (N=13), “insectivore” (N=54), “insectivore/frugivore” (N=37), “insectivore/granivore” (N=27), “nectarivore/insectivore” (N=5), and “omnivore” (N=26) and were assigned using food and feeding descriptions from the Handbook of the Birds of the World (del Hoyo et al. 2017). Omnivores were assigned as species with descriptions including the term “omnivorous”. For further analyses where diet category was included as a predictor, strict frugivores (N=3) and nectarivores/insectivores (N=4) were excluded due to low sample sizes.

Because discrete assigned diet categories may not adequately capture relevant degrees of variation among proportions of food types within avian diets, we also collected published quantitative measurements of species’ diets for N=96 species. We collected percent diet composition for adults, mainly measured as percent stomach content volume (N=81), percent fecal content volume (N=8), or stomach or fecal content frequency proportion (N=7). We separated major food sources into three major categories: Animal, Fruit, and Seed. Finally, we followed (Olson et al. 2008) and used a 7point scoring system to bin Animal, Seed, and Fruit diet proportions for all species. Scoring was necessary to reduce the potential measurement error due to between-study differences in diet categorization, and to include verbal descriptions of species’ diets in cases where precise numerical estimates were not provided. Scores for Animal, Fruit and Seed diet were assigned by the percent contributed to species’ entire diets: 0-1% scored as 1, 1.1-16% scored as 2, 16.8-33% scored as 3, 33.4-50% scored as 4, 50.1-66.7% scored as 5, 66.8-83.4% scored as 6, 83.5-100% scored as 7. For each of these 96 species, our dataset again included: weighted average egg rejection rate (weighted by study sample sizes), average body mass (g), average wing chord (mm), average tarsus length (mm), average length of bill culmen (mm), average bill width and depth (mm) (from Ricklefs 2017), animal diet, seed diet, fruit diet, forage zone, associated brood parasite taxon, and whether the species is a known current host of a brood parasite. We include both species considered to be current hosts or “non-hosts” (*sensu* Medina et al. 2017) in all analyses together because the focal interest of this study is to examine species’ ability to recognize and reject foreign eggs from the nest.

### Comparative Phylogenetic Analyses

For the full dataset, we ran five separate Bayesian MCMCglmm models (Hadfield 2010) predicting egg rejection rates for N=165 species. We ran all MCMCglmm models over 100 phylogenies using the mulTree package (Guillerme & Healy 2018) in R (R Core Team 2017), using a weakly informative parameter expanded prior (V=1, nu=1, prior mean alpha.mu=0, alpha.V=103) (de Villemereuil 2012; Hadfield 2012), setting the number of MCMC generations to 4,000,000, the thinning interval to 1500, and the burn in period to 100,000. Models were run in parallel over 7 chains to obtain at ≥2000 samples per chain. Convergence between model chains was assessed using the Gelman-Rubin statistic, the potential scale reduction factor (PSR), and all were models required to have a PSR below 1.1 (Healy et al. 2014).

We ran five separate models with the full dataset (N=165): 1) testing the influence of species’ foraging zone (aerial, arboreal or ground), 2) testing for differences among associated brood parasite taxon and host status (*Cuculidae* vs. *Molothrus*, current host vs. non-host, and their interaction), 3) testing the influence of species’ categorized diet phenotypes with insectivores set as the comparison group (i.e., insectivore/frugivore, insectivore/granivore, granivore, and omnivore), and two separate models testing the influence of categorized diet phenotypes in 4) current *Molothrus* hosts only (N=56), and 5) current *Cuculidae* hosts only (N=53). Log10-transformed body mass was also included as a predictor in all models to account for the influence of species’ body sizes. For the current hosts only diet category models, strict granivores were excluded due to low samples sizes in the separate samples of *Molothrus* (N=3 granivores) and *Cuculidae* hosts (no granivores).

For the reduced dataset of N=96 species, we first performed two separate phylogenetic principal component analyses (pPCA) (Revell 2009), 1) morphology-based pPCA on log10transformed measurements of body mass, bill length, wing chord, tarsus length, bill length, bill width and bill depth. The first three components explained 87% of all variation in species’ bill and body morphologies, and we named these components based on interpreted axes of their loadings: PC1-Size, PC2-Tarsus vs. Bill Width, and PC3-Bill Length (Table 1).

**Table 1:** *Principal component scores from phylogenetic PCA on morphological measurements of N=96 species*

|  | PC1-Size | PC2-Tarsus vs. Bill Shape | PC3-Bill Length | PC4 | PC5 | PC6 |
| --- | --- | --- | --- | --- | --- | --- |
| Bill length | **-0.76** | 0.24 | **0.56** | -0.21 | -0.03 | 0.04 |
| Bill width | **-0.74** | **-0.54** | 0.09 | 0.17 | -0.35 | -0.01 |
| Bill depth | **-0.82** | -0.33 | 0.05 | 0.17 | 0.43 | 0.04 |
| Body mass | **-0.94** | 0.12 | -0.16 | -0.13 | 0.03 | -0.25 |
| Wing chord | **-0.86** | -0.01 | -0.36 | -0.33 | -0.05 | 0.16 |
| Tarsus length | **-0.74** | **0.52** | -0.11 | 0.4 | -0.08 | 0.06 |
| Standard deviation | 1.99 | 0.86 | 0.7 | 0.63 | 0.57 | 0.3 |
| Proportion of variance | 0.66 | 0.12 | 0.08 | 0.07 | 0.05 | 0.02 |
| Cumulative proportion | 0.66 | 0.78 | 0.87 | 0.93 | 0.98 | 1 |
| PCA model $\lambda$ | 0.87 | | | | | |

Then, we ran a separate diet-based pPCA using Animal, Seed and Fruit diet scores. We included all three components from this pPCA and named them: PC1Animal vs. Seed diet, PC2-Fruit vs. Seed diet, and PC3-Omnivory (Table 2). Phylogenetic signal was high for both the morphology (Pagel’s *λ*=0.87) and diet pPCA (Pagel’s *λ*=0.85). We ran a single MCMCglmm model predicting species’ egg rejection rates with predictors of PC1-Size, PC2-Tarsus vs. Bill Width, PC3-Bill Length, PC1-Animal vs. Seed, PC2-Fruit vs. Seed and PC3-Omnivory over 100 phylogenies using the same model prior and parameters as described above. Egg rejection rates were mean-centered and scaled to one standard deviation from the mean for all models.

**Table 2:** *Principal component scores from phylogenetic PCA on animal, fruit and seed diet scores of N=96 species*

|  | PC1-Animal vs. Seed | PC2-Fruit vs. Seed | PC3-Omnivory |
| --- | --- | --- | --- |
| Animals | **0.98** | 0.16 | 0.11 |
| Fruits | -0.4 | **-0.9** | 0.15 |
| Seeds | **-0.77** | **0.62** | 0.12 |
| Standard deviation | 0.4 | 0.29 | 0.06 |
| Proportion of variance | 0.64 | 0.34 | 0.02 |
| Cumulative proportion | 0.64 | 0.98 | 1 |
| PCA model $\lambda$ | 0.85 | | |

## III. RESULTS

All MCMCglmm models passed convergence (PSR ≤ 1.1) and produced ≥ 1500 posterior estimates per each predictor from 100 iterations of each model, where each iteration was run with a separate phylogenetic hypothesis. For each model predictor, we report its posterior mode and 95% highest density interval (HDI), calculated from the highest density region of combined posterior distributions across all 100 model iterations. We also provide estimates of posterior probabilities of effects of predictors having negative (P-), null (P0, probability of predictor having an effect within an interval around zero), and positive (P+) effects on egg rejection rates, calculated using the BayesCombo package in R (Contrino et al. 2017).

### Categorized diet predicting egg rejection

For the full dataset models including all species associated with both *Molothrus* and *Cuculidae* brood parasites and both current hosts and non-hosts, omnivorous (posterior mode=0.45, 95%-HDI=[0,0.90], P-|P0|P+= 0.04|0.2|0.77) and insectivore/frugivore species (posterior mode=0.44, 95%-HDI=[0.05,0.84], P-|P0|P+= 0.03|0.15|0.82) have higher egg rejection rates in comparison with mainly insectivorous species, while granivorous species have relatively lower egg rejection rates (granivore posterior mode=-0.63, 95%-HDI=[-1.30,0.05], P-|P0|P+= 0.75|0.21|0.04; granivore/insectivore posterior mode=-0.49, 95%HDI=[ −0.93, −0.06], P-|P0|P+= 0.84|0.13|0.02) (Fig.1). Higher body mass is associated with higher egg rejection rates (Log10-Body Mass posterior mode=0.12, 95%-HDI=[-0.08,0.32], P-|P0|P+= 0.1|0.35|0.55). Residual variance posterior mode=0.35, 95%-HDI=[0.16,0.59]. Phylogenetic variance posterior mode=0.90, 95%-HDI=[0.20,1.95].

### Forage zone predicting egg rejection

In comparison to ground foraging species, arboreal foragers have marginally higher egg rejection rates (posterior mode=0.21, 95%-HDI=[-0.14,0.56], P-|P0|P+= 0.11|0.36|0.53), whereas no discernable pattern was found in the comparison between aerial and ground foragers (posterior mode=0.09, 95%-HDI=[-0.65,0.79], P-|P0|P+= 0.25|0.42|0.33) (Fig. 1). In concordance with the diet category models, higher body mass is associated with higher egg rejection rates (Log10-Body Mass posterior mode=0.18, 95%-HDI=[-0.01,0.39 P-|P0|P+= 0.05|0.23|0.72) (Fig.1). Residual variance posterior mode=0.27, 95%-HDI=[0.11,0.52]. Phylogenetic variance posterior mode=1.64, 95%HDI=[0.71,2.74].

**Figure 1:**
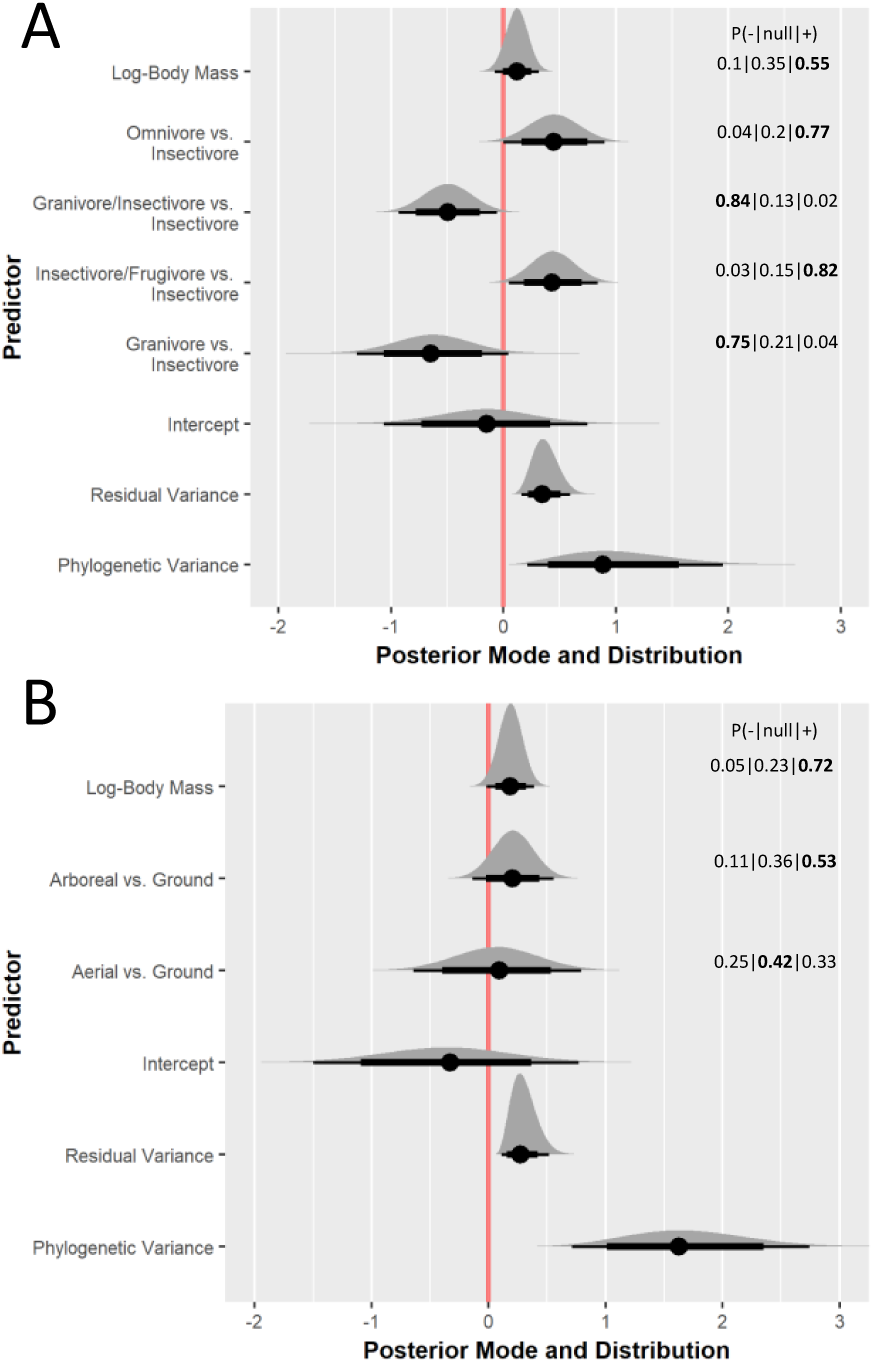
*Full dataset models testing the influence of* ***A*** *diet and* ***B*** *forage zone on egg rejection rates in N=165 species. Posterior modes and distributions, their 95% highest-density intervals (thin line), 80% highest-density intervals (thick line), and posterior probabilities of having either negative, null or positive effects on egg rejection rates are presented*.

### Associated brood parasite taxon and host status predicting egg rejection

Species associated with *Cuculidae* brood parasites have lower egg rejection rates than species associated with *Molothrus* brood parasites (posterior mode=-0.86, 95%-HDI=[-1.44,0.28], P-|P0|P+= 0.97|0.03|0), and current host species have lower egg rejection rates than non-host species (posterior mode=-0.46, 95%HDI=[-0.90,-0.01], P-|P0|P+= 0.79|0.18|0.03) (Fig.2). Accordingly, because both associated brood parasite taxon and host/nonhost status combined predict a substantial amount of variation in species’ egg rejection rates (interaction between associated brood parasite taxon and host status posterior mode=1.18, 95%-HDI=[0.55,1.80], P-|P0|P+= 0|0|1), we ran separate diet category models predicting egg rejection rates of current hosts of *Cuculidae* brood parasites and current hosts of *Molothrus* brood parasites. Residual variance posterior mode=0.28, 95%-HDI=[0.12,0.48]. Phylogenetic variance posterior mode=1.42, 95%HDI=[0.67,2.46].

**Figure 2:**
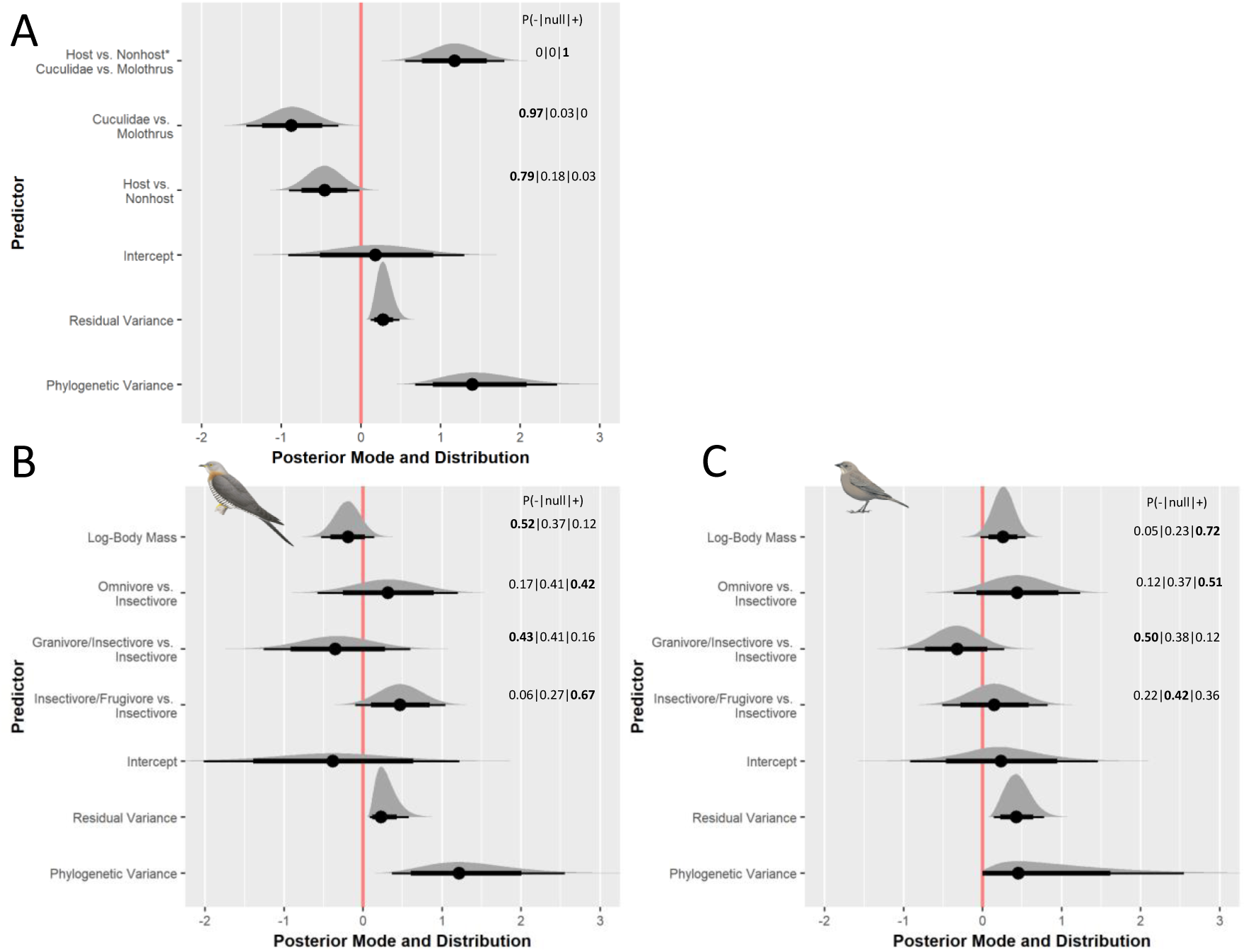
*Full dataset model testing the influence of* ***A*** *associated brood parasite taxon and host status in N=165 species. Reduced dataset models testing the influence of diet on egg rejection rates in current hosts of* ***B*** *Cuculidae (N=53) and* ***C*** *Molothrus (N=56) brood parasites. Posterior modes and distributions, their 95% highest-density intervals (thin line), 80% highest-density intervals (thick line), and posterior probabilities of having either negative, null or positive effects on egg rejection rates are presented*.

### Categorized diet predicting egg rejection in ***Cuculidae* and *Molothrus* hosts**

In *Cuculidae* hosts (N=53 species), larger body mass is marginally associated with lower egg rejection rates (Log10-Body Mass posterior mode=-0.19, 95%-HDI=[-0.53,0.02], P-|P0|P+= 0.52|0.37|0.12) (Fig.2). In comparison with mainly insectivorous hosts, omnivores have marginally higher egg rejection rates (posterior mode=0.32, 95%-HDI=[-0.57,1.20], P-|P0|P+= 0.17|0.41|0.42) and insectivore/frugivores have credibly higher egg rejection rates (posterior mode=0.47, 95%HDI=[-0.10, 1.04], P-|P0|P+= 0.06|0.27|0.67). Granivore/insectivore *Cuculidae* hosts have marginally lower egg rejection rates than insectivore hosts (posterior mode=-0.33, 95%HDI=[-1.26,0.60], P-|P0|P+= 0.43|0.41|0.16). Residual variance posterior mode=0.23, 95%HDI=[0.08,0.58]. Phylogenetic variance posterior mode=1.21, 95%-HDI=[0.36,2.55].

In *Molothrus* hosts (N=56 species), larger body mass is associated with higher egg rejection rates (Log10-Body Mass posterior mode=0.26, 95%-HDI=[-0.02,0.54], P-|P0|P+= 0.05|0.23|0.72) (Fig.2). In comparison with mainly insectivorous hosts, omnivores have marginally higher egg rejection rates (posterior mode=0.44, 95%-HDI=[-0.37,1.24], P-|P0|P+= 0.12|0.37|0.51) but insectivore/frugivore diet type does not credibly predict egg rejection rates (posterior mode=0.15, 95%HDI=[-0.51,0.82], P-|P0|P+= 0.22|0.42|0.36). Granivore/insectivore *Molothrus* hosts have marginally lower egg rejection rates than insectivore hosts (posterior mode=-0.33, 95%HDI=[-0.95,0.27], P-|P0|P+= 0.50|0.38|0.12). Residual variance posterior mode=0.42, 95%HDI=[0.14,0.78]. Phylogenetic variance posterior mode=0.46, 95%-HDI=[-0.02,2.54].

### Quantitative diet and egg rejection

In a subset of N=96 species for which we collected quantitative measures of diet, we found PC3-Omnivory to be negatively associated with egg rejection rates (posterior mode=-0.39, 95%HDI=[-0.87,0.10], P-|P0|P+= 0.65|0.28|0.07), greater frugivory (PC2-Fruit vs. Seed*-1 posterior mode=0.20, 95%-HDI=[-0.31,-0.10], P- |P0|P+= 0|0|1) and consumption of animals (PC1-Animal vs. Seed posterior mode=0.08, 95%-HDI=[0,0.16], P-|P0|P+= 0.03|0.18|0.78) to be positively associated with higher egg rejection rates (Fig.3). For morphological predictors, larger species have higher egg rejection rates (PC1-Size*-1 posterior mode=0.02, 95%-HDI=[0.03, 0], P-|P0|P+= 0.01|0.06|0.93) but PC2-Tarsus vs. Bill Width was not predictive of species’ egg rejection rates (posterior mode=0.01, 95%-HDI=[-0.02,0.05], P-|P0|P+= 0.19|0.42|0.39). Bill length, as captured by our pPCA, seems to have no direct association with egg rejection rates (PC3-Bill Length posterior mode=0, 95%-HDI=[-0.05,0.05], P-|P0|P+= 0.3|0.42|0.29). Residual variance posterior mode=0.38, 95%-HDI=[0.18,0.66]. Phylogenetic variance posterior mode=0.64, 95%HDI=[0.01,1.73].

**Figure 3:**
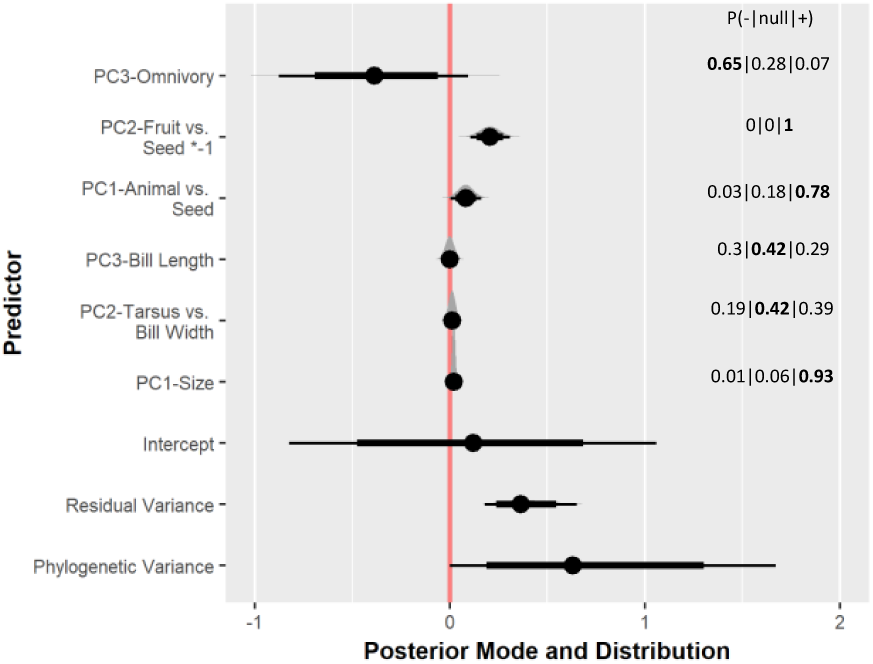
Reduced dataset model testing the effect of quantitatively measured diets on egg rejection rates in N=96 species. Diet data collected from published studies where physical contents of consumed items were observed (i.e., stomach or fecal content). Posterior modes and distributions, their 95% highest-density intervals (thin line), 80% highest-density intervals (thick line), and posterior probabilities of having either negative, null or positive effects on egg rejection rates are presented.

## IV. DISCUSSION

Across all species sampled, omnivorous and frugivorous diet rather than insectivorous diet, arboreal rather than ground foraging, and relatively large body mass are predictive of greater ability to recognize and reject foreign eggs (Fig.1). When accounting for the influence of host-brood parasite coevolutionary relationships, directional effects of all diet types on egg rejection rates are similar across hosts of both *Cuculidae* and *Molothrus* hosts (Fig.2), albeit to a less pronounced degree than when testing effects of diets on egg rejection in both current hosts and non-hosts associated with both brood parasite taxa (Fig.1). Furthermore, these patterns between diet types and egg rejection rates remain consistent when diets are measured as quantitative, rather than categorical traits (Fig.3)—except for our diet principal component score interpreted as characterizing an omnivorous diet (i.e., PC3-Omnivory, which explains 2% of the variation in diet in the reduced N=96 dataset, shown to have a negative effect on egg rejection–Table 2), which may not have adequately captured omnivory.

An alternative explanation for the patterns we found between diet types and egg rejection rates may be that species with diets considered to be unsuitable for brood parasites are far less likely to be parasitized, and therefore do not need to evolve egg rejection defenses. Specifically, diet unsuitability may explain the lack of egg rejection defenses we found in highly granivorous species (Fig.1,2). Indeed, a recent study found that common cuckoos Cuculus canorus prefer to parasitize host species that feed their nestlings insects over those that do not (Stokke et al. 2018), and *Molothrus* cowbird nestlings are known to require animal protein to survive and fledge from host nests (Mason 1986). However, many potential host species have highly insectivorous diets during their breeding season and switch to eating mainly seeds and grains in the nonbreeding season, and we account for these species in our analyses (i.e. “granivore/insectivore”). We also caution that adults’ diets may not directly correspond to the diets which they feed to their nestlings. For example, adult starlings (*Sturnus vulgaris*) consume large amounts of fruit, but their nestlings do not, even when fruits are available during the nestling feeding period (Moeed 1980). Because our data are drawn from descriptions and data available for adults only, they likely do not provide an accurate representation of diets fed to nestlings.

Our results suggest phenotype traits associated with different foraging strategies and diets (e.g., sensory systems and morphology), in combination with the degree of brood parasite egg mimicry (Stoddard & Stevens 2011) and potential costs associated with being parasitized (Medina & Langmore 2015), may constrain or facilitate the evolution of host species’ ability to recognize and reject a foreign egg by either its color (Stoddard & Hauber 2017), maculation pattern, or shape and size. For example, visually-guided insectivorous birds possess visual fields predicted to be better-suited for eye-beak coordination than birds that consume immobile food (Moore et al. 2015), and better eye-beak coordination may confer greater ability to physically remove a foreign egg from the nest.

Additionally, frugivory is facilitated by use of chromatic cues and contrasts (Cazetta et al. 2009) and the range of fruits which can be consumed is limited by minimum bill gape-width (Wheelwright 1985), which may also carry over into discrimination between colors of own vs. foreign eggs and ability to grasp and remove an egg from the nest (Rohwer & Spaw 1988). Because body mass and eye size are highly positively correlated in birds (Garamszegi et al. 2002), and larger eye size is associated with higher visual acuity (Kiltie 2000), the positive relationships we found between body mass and egg rejection rates may indicate that greater visual acuity, along with larger body and bill size, confers a greater ability to recognize and reject foreign eggs. Whereas we found evidence that certain diet and foraging ecology-related phenotypes influence foreign egg recognition and rejection rates, our study does not directly examine specific sensory and physical mechanisms that may influence egg rejection defenses. Therefore, we suggest future comparative and experimental studies use our results as a guideline for examining suites of morphological, sensory or cognitive traits that may form a mechanism explaining the patterns we found here (e.g., complete bill morphometrics, visual acuity, photoreceptor densities, color discrimination ability, etc.).

In summary, we provide exploratory support for the hypothesis that foraging ecology and diet may affect the evolutionary trajectory of egg rejection defenses in avian brood parasite hosts. Specifically, we predict current and future hosts of avian brood parasites that do not currently exhibit egg rejection defenses to more readily evolve egg rejection if they consume mainly animals and fruits rather than seeds and grains, forage arboreally rather than in the air or on the ground and have a relatively large body size.

## ACKNOWLEDGEMENTS

We thank M.M. Louder, M. Abolins-Abols, and E. Fernandez-Juricic for their help to improve the manuscript.

## AUTHOR CONTRIBUTIONS

AB Luro and ME Hauber conceived the study, AB Luro collected and analyzed the data, and both authors wrote and edited the manuscript.

---

**Figure S1:**
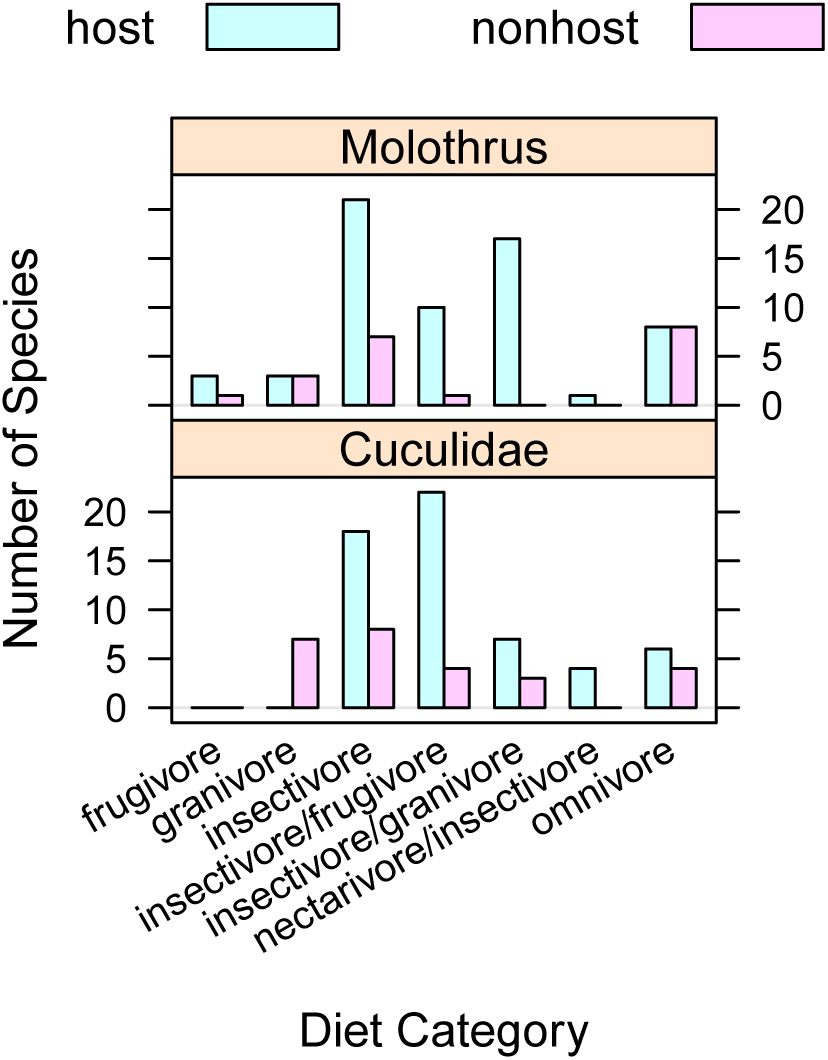
Number of species included in dataset by diet category, associated brood parasite taxon and current host (blue)/non-host (pink) status.

**Figure S2:**
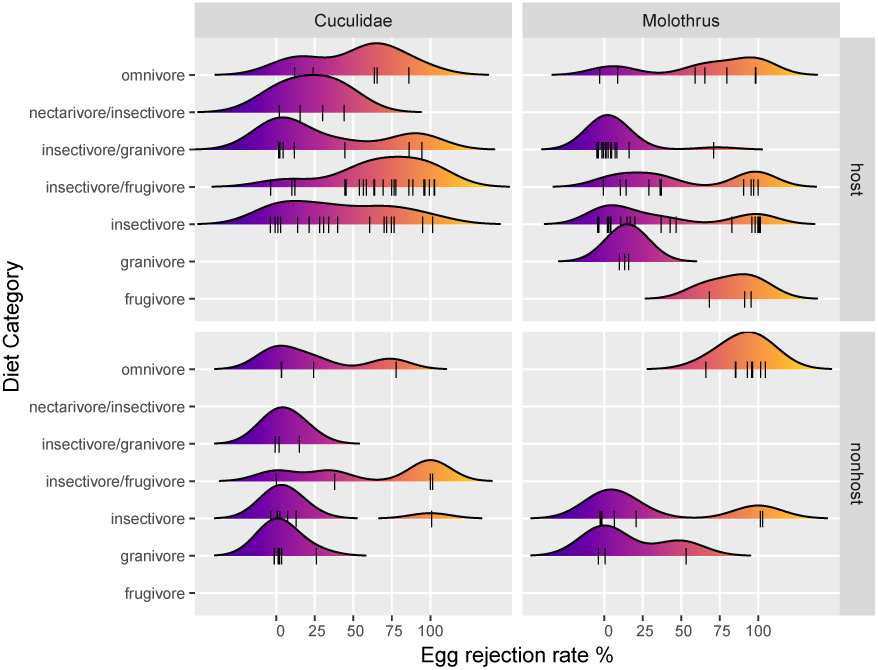
Density distributions of egg rejection rates within each diet category, grouped by associated brood parasite taxon (columns) and current host/nonhost status (rows). Each vertical tick represents an observation from a single species.

**Figure S3:**
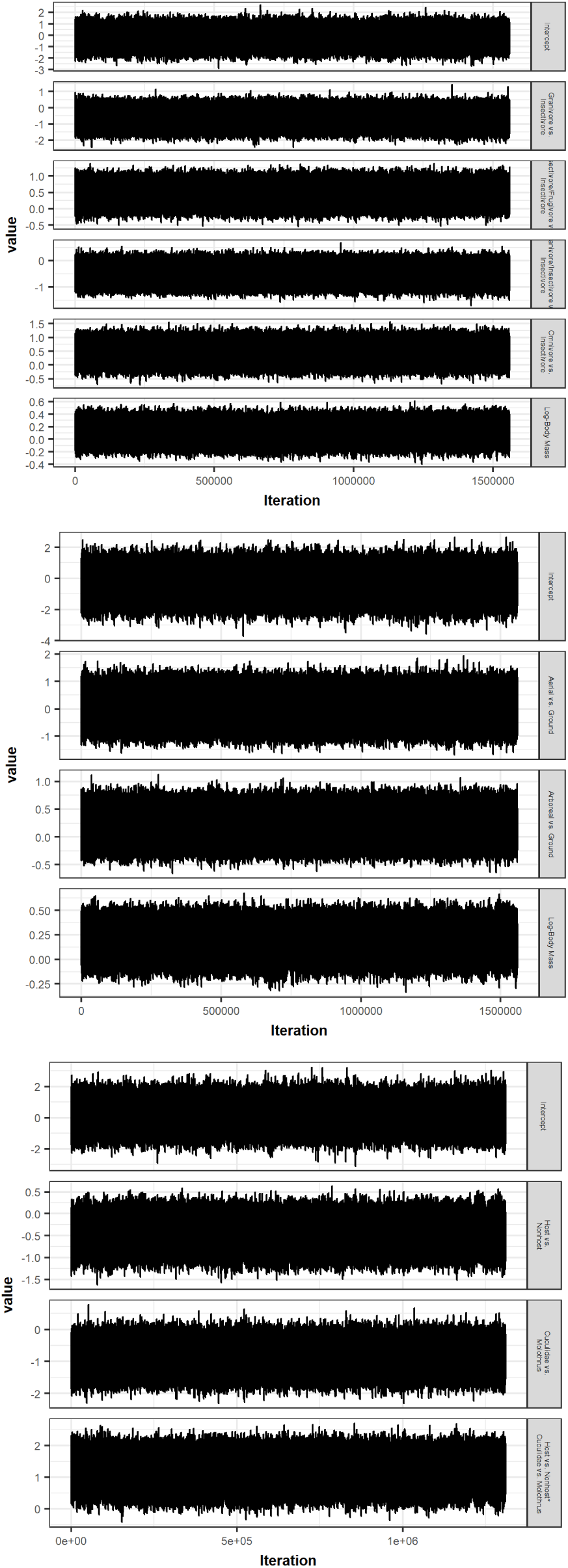
Traceplots from MCMCglmm models where full dataset was used. From top to bottom: Categorized diet model traces, Forage zone model traces, Associated brood parasite taxon and host status model traces.

**Figure S4:**
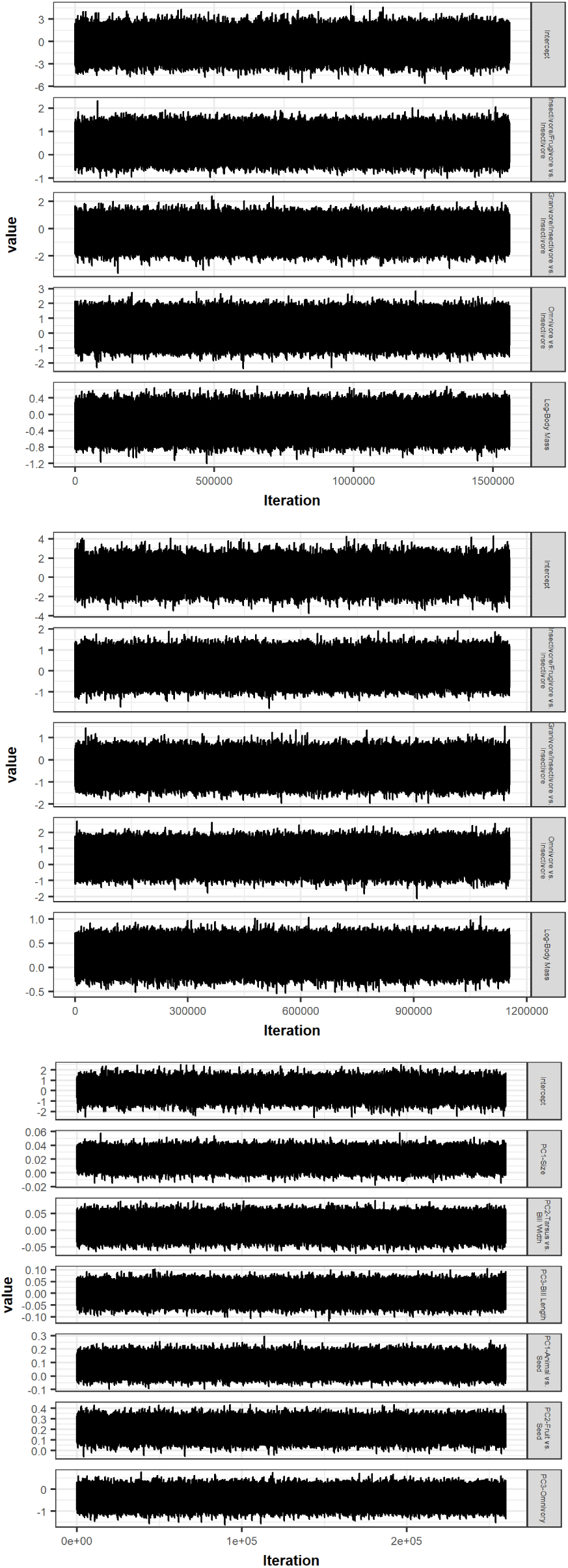
Traceplots from MCMCglmm models where reduced datasets were used. From top to bottom: Cuculidae categorized diet model traces, Molothrus categorized diet model traces, Quantitative diet model traces.

